# Detecting subtle transcriptomic perturbations induced by lncRNAs Knock-Down in single-cell CRISPRi screening using a new sparse supervised autoencoder neural network

**DOI:** 10.1101/2023.07.11.548494

**Authors:** Marin Truchi, Caroline Lacoux, Cyprien Gille, Julien Fassy, Virginie Magnone, Rafael Lopez-Goncalvez, Cédric Girard-Riboulleau, Iris Manosalva-Pena, Marine Gautier-Isola, Kevin Lebrigand, Pascal Barbry, Salvatore Spicuglia, Georges Vassaux, Roger Rezzonico, Michel Barlaud, Bernard Mari

**Affiliations:** Université Côte d’Azur, IPMC, CNRS UMR7275, IHU RespiERA, Valbonne, France; Université Côte d’Azur, I3S, CNRS UMR7271, Sophia Antipolis, France; Université Aix-Marseille, Inserm, TAGC, UMR1090, Marseille, France

## Abstract

Single-cell CRISPR-based transcriptome screens are potent genetic tools for concomitantly assessing the expression profiles of cells targeted by a set of guides RNA (gRNA), and inferring target gene functions from the observed perturbations. However, due to various limitations, this approach lacks sensitivity in detecting weak perturbations and is essentially reliable when studying master regulators such as transcription factors. To overcome the challenge of detecting subtle gRNA induced transcriptomic perturbations and classifying the most responsive cells, we developed a new supervised autoencoder neural network method. Our Sparse supervised autoencoder (SSAE) neural network provides selection of both relevant features (genes) and actual perturbed cells. We applied this method on an in-house single-cell CRISPR-interference-based (CRISPRi) transcriptome screening (CROP-Seq) focusing on a subset of long non-coding RNAs (lncRNAs) regulated by hypoxia, a condition that promote tumor aggressiveness and drug resistance, in the context of lung adenocarcinoma (LUAD). The CROP-seq library of validated gRNA against a subset of lncRNAs and, as positive controls, HIF1A and HIF2A, the 2 main transcription factors of the hypoxic response, was transduced in A549 LUAD cells cultured in normoxia or exposed to hypoxic conditions during 3, 6 or 24 hours. We first validated the SSAE approach on HIF1A and HIF2 by confirming the specific effect of their knock-down during the temporal switch of the hypoxic response. Next, the SSAE method was able to detect stable short hypoxia-dependent transcriptomic signatures induced by the knock-down of some lncRNAs candidates, outperforming previously published machine learning approaches. This proof of concept demonstrates the relevance of the SSAE approach for deciphering weak perturbations in single-cell transcriptomic data readout as part of CRISPR-based screening.

## Introduction

Cancer cells in solid tumors often suffer from hypoxic stress and adapt to this micro-environment via the activation of Hypoxia inducible factor (HIF), a heterodimeric transcription factor composed of either HIF-1*α* or HIF-2*α* (initially identified as endothelial PAS domain protein (EPAS1)) and HIF-1*β* /ARNT subunits [1–3]. In normoxia, HIF*α* is continuously degraded by an ubiquitin–dependent mechanism mediated by interaction with to the von Hippel–Lindau (VHL) protein. Hydroxylation of proline residues in HIF*α* is necessary for VHL binding and is catalyzed by the *α*-ketoglutarate-dependent dioxygenases prolyl hydroxylases (PHD). During hypoxia, PHDs are inactive, leading to HIF-*α* stabilization, dimerization with HIF-1*β* and finally translocation into the nucleus to bind to E-box-like hypoxia response elements (HREs) within the promoter region of a wide range of genes that control cellular oxygen homeostasis, erythrocyte production, angiogenesis and mitochondrial metabolism [4]. These molecular changes are notably crucial for cells to adapt to stress by lowering oxygen consumption by shifting from oxidative metabolism to glycolysis. While HIF-1 and HIF-2 bind to the same HRE consensus sequence, they are non-redundant and have distinct target genes and mechanisms of regulation. It is generally accepted that the individual HIFs have specific temporal and functional roles during hypoxia, known as the HIF switch, with HIF-1 driving the initial response and HIF-2 directing the chronic response [5]. In most solid tumors, including lung adenocarcinoma (LUAD), the degree of hypoxia is associated with poor clinical outcome. Induction of HIF activity upregulates genes involved in many hallmarks of cancer, including metabolic reprogramming, epithelial-mesenchymal transition (EMT), invasion and metastasis, apoptosis, genetic instability and resistance to therapies. Emerging evidence have highlighted that hypoxia regulates expression of a wide number of non-coding RNAs classes including microRNAs (miRNAs) and long non-coding RNAs (lncRNAs) that in turn are able to influence the HIF-mediated response to hypoxia [6–8]. LncRNAs constitute a heterogeneous class of transcripts which are more than 200 nt long with low or no protein coding potential, such as intergenic and antisense RNAs, transcribed ultraconserved regions (T-UCR) as well as pseudogenes. Recent advances in cancer genomics have highlighted aberrant expression of a wide set of lncRNAs [9], revealing their roles in regulating the genome at several levels, including genomic imprinting, chromatin state, transcription activation or repression, splicing and translation control [10]. LncRNAs can regulate gene expression through different mechanisms, as guide, decoy, scaffold, miRNA sponges or micropeptides. Of note, recent studies demonstrated the role of several lncRNAs in the direct and indirect regulation of HIF expression and pathway through diverse mechanisms [7]. Moreover, hypoxia-responsive lncRNAs have been shown to play regulatory functions in pathways associated with the hallmarks of cancer. For instance, the hypoxia-induced Nuclear-Enriched Abundant Transcript 1 (NEAT1) lncRNA has been associated with the formation of nuclear structures called paraspeckles during hypoxia as well as an increased clonogenic survival of breast cancer cells. Another highly studied lncRNA, Metastasis-Associated Lung Adenocarcinoma Transcript 1 (MALAT1, also known as NEAT2) has been found upregulated by hypoxia in LUAD A549 cells and associated with various cellular functions depending on tumor cell types including cell death, proliferation, migration and invasion [11]. Starting from an expression screening in LUAD patients samples and cell lines subjected to hypoxia, we have characterized a new nuclear hypoxia-regulated transcript from the Lung Cancer Associated Transcript (LUCAT1) locus associated with patient prognosis and involved in redox signaling with implication for drug resistance [12]. Additional promising lncRNAs candidates regulated by hypoxia and/or associated with bad prognosis have been identified but deciphering the regulatory functions of these poorly annotated transcripts remains a major challenge. Pooled screening approaches using CRISPR-based technology have offered the possibility to evaluate mammalian gene function, including lncRNAs at genome scale levels [13]. More recently, they have been applied to cancer cell lines and have confirmed the oncogenic or tumor suppressor roles of some lncRNAs [14]. This strategy is able to test a large number of candidates simultaneously but require well identified phenotypes such as cell proliferation, cell viability, or cell migration. More subtle screens require techniques based on transcriptomic signatures [15] and approaches have been developed to combine CRISPR gene manipulation, including CRISPR interference and single-cell RNA-seq (scRNA-seq) based on droplet isolation, such as Perturb-seq [16], CROP-seq [17] and ECCITE-seq [18]. These methods combine the advantages of screening a large number of genes simultaneously and linking the modifications to the transcriptomic phenotype, all by breaking down the perturbation signal cell by cell [10, 16].

In single cell omics applications, most of the quantified features are weakly detected, resulting in large, sparse and noisy data which required feature selection to extract biologically relevant signals [19]. Moreover, cells are often grouped according to their phenotype and/or their experimental condition in order to compare features quantification between the defined cell classes. However, the intra-classes heterogeneity can mask a signal of interest. This is particularly the case in the context of CRISPRi screens with a single-cell transcriptomic readout where the inhibition level of the target gene varies between each cell and induces a more or less detectable perturbation signature. Classification tools such as Mixscape [20], based on Mixture Discriminant Analysis [21], has proven efficacy to identify strong CRISPR-induced effects but was unable to detect subtle weak transcriptomic perturbations.

In the present work, we have developed a single-cell CRISPR-interference-based (CRISPRi) transcriptome screening based on the CROP-Seq approach to gain insight on the regulatory functions of hypoxia-regulated lncRNAs. As a proof-of-concept, we generated a CROP-seq library, including validated guide RNAs (gRNA) targeting six previously identified lncRNAs regulated by hypoxia and/or associated with bad prognosis [12] as well as the two master transcription factors of the hypoxic response (HIF1A and HIF2/EPAS1) and negative control guides. To optimize analysis of fine-tuned regulations in this dataset, we have adapted a Sparse supervised autoencoder (SSAE) neural network [22], where we relax the parametric distribution assumption of classical VAE. It leverages on the known cell labels, corresponding to the received gRNA, and a classification loss to incite the latent space to fit the true data distribution. We first validated the approach on HIF1 and HIF2/EPAS1 knock-down, showing a good sensitivity to detect the known temporal switch between both regulators. We then applied the SSAE to the cells treated with the different hypoxia-regulated lncRNAs gRNA to identify subtle signatures linked to the knock-down of the lncRNAs.

## Materials and methods

### Lentivirus production

Lentiviruses were produced using a standard Lipofectamine 2000™transfection protocol, using one million HEK293 cells seeded in a 25 cm2 flask in DMEM medium supplemented with 10% bovine serum. A mixture of four plasmids (3 μg pMDLg/pRRE (addgene “12251”), 1.4 μg pRSV-Rev (addgene “12253”), 2 μg pVSV-G (addgene “12259”) and 2.5 μg of the plasmid containing the expression cassette to package the pooled CROP-seq guides) was transfected. Forty-eight hours later, the medium was collected, centrifuged for 5 minutes at 3000 rpm, and 2.5 mL supernatant containing the viral particles was collected and used to infect cells or aliquoted and stored at -80°C. Large scale preparations of lentivirus were produced at the Vectorology facility, PVM, Biocampus (CNRS UMS3426), Montpellier, France.

### Generation of dCas9-expressing A549 cell line

The lung adenocarcinoma cell line A549 was infected with a lentivirus produced from the plasmid lenti-dCas9-KRAB-MeCP2 (a gift from Andrea Califano, addgene 122205) allowing the expression of a fusion protein MeCP2-KRAB-dCas9 and a gene conferring resistance to blasticidin. Infected cells were then grown in the presence of 10 μg/mL of blasticidin (Sigma). Selection of A549-KRAB-MeCP2 cells was complete within 3 to 5 days. Bulk blasticidin positive cells were amplified and cloned for the CRISPRi scRNA-seq experiments. The best clone was selected according to the expression level of MeCP2-KRAB-dCas9 mRNA and to the most effective inhibition of NLUCAT1 using the NLUCAT1 sg3 RNA.

### Cloning of individual guides in the CROPseq-Guide-Puro plasmid

The plasmid CROPseq-Guide-Puro (Datlinger et al. Nat Methods 2017) (a gift from C Bock, Addgene plasmid 86708) was digested using the restriction enzyme BsmBI (NEB R0580) for 2h at 50°C. The relevant fragments (around 8 kB) were gel-purified using the Qiagen Gel purification kit and stored at –20°C in 20-fmol aliquots. Guides against the targeted genes (see Supplemental Table 1 for selected sequences) were cloned using the Gibson assembly method (NEBuilder HiFi DNA Assembly Master Mix, NEB E2621). Aliquoted, BsmBI-digested plasmid was mixed with 0.55 μL guide oligonucleotide (200nM) in 10μl total volume, combined with 10μl 2X NEBuilder HiFi Assembling Master mix and the mixture was incubated at 50°C for 20 minutes. 8μL of NEBuilder Assembling mixture were incubated with 100 μL of Stabl2 competent E coli. The mixture was heat-shocked at 42°C for 45 seconds and transferred to ice for 2 minutes. SOC medium (900 μl) was added to the Stabl2-NEBuilder mixture and the mix was incubated at 37°C for 1 hour. Transformed bacterial cells (350μl) were plated onto LB agarose plates containing ampicillin (100μg/mL) and incubated overnight at 37°C. Individual colonies were picked and grown overnight in 5 mL of Terrific Broth medium containing 150μg/mL ampicillin and low-endotoxin, small scale preparation of plasmid DNA were performed using the ToxOut EndoFree Plasmid Mini Kit from BioVision (K1326-250). All plasmids were verified by Sanger sequencing with the primer 5’-TTGGGCACTGACAATTCCGT-3’.

### Selection of the guides

A549-KRAB-MeCP2 cells were infected with lentivirus obtained from individual CROPseq-Guide-Puro plasmids, encoding individual guides. Infected cells were then grown in the presence of 1 μg/mL of puromycin (Sigma). A week later, total RNAs were purified from A549-KRAB-MeCP2 cells infected with guide encoding lentiviruses and RT-qPCR (primers sequences presented in Supplemental Table 2) were performed to measure expression of the targeted genes. A validated guide was defined as a guide providing at least 75% inhibition of targeted gene expression compared to a control guide.

### Lentiviral transduction with gRNA libraries and cell preparation for chromium scRNA-seq

A549-KRAB-MeCP2 cells were transduced with different amounts of the viral stock containing the library of pooled, selected gRNA. After six hours, the virus-containing medium was replaced by fresh complete culture medium. Puromycin selection (1μg/ml) was started at 48 h post-transduction, and two days later, the plate with about 30% surviving cells was selected, corresponding roughly to a MOI=0.3. The cells were then amplified under puromycin selection for 5 days. The cells were then plated and further cultured in normoxia or in hypoxic condition (1% O2) for 3h, 6 h or 24h. Cells were trypsinized counted and assessed for cell viability using the Countess 3 FL (Fisher Scientific). Samples were then stained for multiplexing using cell hashing [23], using the Cell Hashing Total-Seq-ATM protocol (Biolegend) following the protocol provided by the supplier, using 4 distinct Hash Tag Oligonucleotides-conjugated mAbs (TotalSeq™-B0255, B0256, B0257 and B0258). Briefly, for each condition, 1.106 cells were resuspended in 100μL of PBS, 2% BSA, 0.01% Tween and incubated with 10μL Fc Blocking reagent for 10 minutes at 4°C then stained with 0.5μg of cell hashing antibody for 20 minutes at 4°C. After washing with PBS, 2% BSA, 0.01% Tween, samples were counted and merged at the same proportion, spun 5 minutes 350 x g at 4°C and resuspended in PBS supplemented with 0.04% of bovine serum albumin at final concentration of 500 cells/μL. Samples were then adjusted to the same concentration, mixed in PBS supplemented with 0.04% of bovine serum albumin at a final concentration of 100 cells/μl and pooled sample were immediately loaded onto10X Genomics Chromium device to perform the single cell capture.

### Generation of CROP-seq librairies and single-cell RNA-seq data processing

After single-cell capture on the 10X Genomics Chromium device (3’ V3), libraries were prepared as recommended, following the Chromium Next GEM Single Cell 3’ Reagent Feature Barcoding V3.1 kit (10X Genomics) and a targeted gRNA amplification [24] with respectively 6, 8 and 10 PCR cycles. Libraries were then quantified, pooled (80% RNA libraries, 10% gRNA libraries and 10% hashing libraries) and sequenced on an Illumina NextSeq 2000. Alignment of reads from the single cell RNA-seq library and unique molecular identifiers (UMI) counting, as well as oligonucleotides tags (HTOs) counting, were performed with 10X Genomics Cell Ranger tool (v3.0.2). Reads of the gRNA library were counted with CITE-seq-Count (v1.4.2). Cells without gRNA counts were discared. Counts matrices of total UMI, HTOs, and gRNA were thus integrated on a single object using Seurat R package (v4.1.0), from which the data were processed for analysis. On the total of 19663 cells, 817 cells without gRNA counts were discared.

HTOs and gRNA were demultiplexed with HTODemux() and MULTIseqDemux(autoThresh = TRUE) functions respectively, in order to assign treatment and received gRNA for each cell. On the remaining 18846 cells, only cells identified as “Singlet” after demultiplexing of HTO counts were conserved (14276 cells). The repartition of cells assigned as “Doublet” (high expression of at list 2 different gRNA), “Negative” (no detected gRNA) and “Singlet” (a unique detected gRNA) in all conditions is showed in (**Table 2**). Finally, after transforming the data of the subset of “Singlet” cells using SCTransform(), computing PCA, and performing KNN clustering, 2 clusters of low UMI content and high mitochondrial content cells (3087 cells) were eliminated for the rest of the analysis.

**Table 1.** gRNA library.

| gRNA | Target | Type | % Inhibition |
| --- | --- | --- | --- |
| HIF1A-sg1 | HIF1A | Hypoxic response regulator | >95% |
| HIF1A-sg2 |  |  | >95% |
| HIF2-sg5 | HIF2/EPAS1 |  | >95% |
| LINC00152-sg3 | CYTOR | Hypoxia-regulated lncRNA | >75% |
| LUCAT-sg3 | LUCAT1 |  | >97% |
| LUCAT-sg5 |  |  | >90% |
| MALAT-sg1 | MALAT1 |  | >95% |
| NEAT1-sg2 | NEAT1 |  | >85% |
| NEAT1-sg6 |  |  | >95% |
| SNHG12-sg1 | SNHG12 |  | >75% |
| SNHG12-sg3 |  |  | >90% |
| SNHG21-sg5 | SNHG21 |  | >85% |
| Neg-sg1 | None | Negative control | None |
| Neg-sg2 |  |  |  |

**Table 2.** Repartition of Doublet, Singlet, and Negative cells in all conditions after demulplexing of gRNA counts.

|  | Normoxia (%) | Hypoxia-3h (%) | Hypoxia-6h (%) | Hypoxia-24h (%) |
| --- | --- | --- | --- | --- |
| Doublet | 8,378 | 9,908 | 7,717 | 8,096 |
| HIF1A-sg1 | 6,752 | 5,766 | 5,681 | 5,181 |
| HIF1A-sg2 | 8,346 | 8,236 | 8,110 | 8,265 |
| HIF2-sg5 | 4,720 | 5,063 | 3,859 | 5,157 |
| LINC00152-sg3 | 4,720 | 4,918 | 5,645 | 5,108 |
| LUCAT1-sg3 | 5,345 | 5,911 | 5,288 | 6,048 |
| LUCAT1-sg5 | 8,690 | 8,842 | 9,218 | 10,120 |
| MALAT1-sg1 | 6,471 | 6,686 | 7,181 | 5,976 |
| NEAT1-sg2 | 5,220 | 5,354 | 4,823 | 5,373 |
| NEAT1-sg6 | 3,501 | 4,288 | 3,930 | 3,952 |
| SNHG12-sg1 | 4,189 | 3,125 | 3,823 | 3,759 |
| SNHG12-sg3 | 2,094 | 2,253 | 1,751 | 1,639 |
| SNHG21-sg5 | 6,690 | 6,492 | 6,967 | 7,398 |
| Neg-sg1 | 6,346 | 5,838 | 6,717 | 6,554 |
| Neg-sg2 | 7,221 | 6,783 | 7,503 | 7,373 |
| Negative | 11,316 | 10,538 | 11,790 | 10,000 |
| Total cell number | 3199 | 4128 | 2799 | 4150 |

### Method: a new sparse supervised autoencoder neural network (SSAE)

#### State of the art of neural networks methods

Deep neural networks have proven their efficiency for classification and feature selection in many domains [25], and have also been applied to omics data analyses [26, 27]. Among the proposed neural networks architectures, autoencoders are able to learn a representation of the data, typically in a latent space of lower dimension than the input space. As such, they are often used for dimensionality reduction [28] and have applications in the medical field as data denoisers or relevant feature selectors [29, 30]. A widely used type of autoencoders is the Variational Autoencoder (VAE) [31]. This VAE adds the assumption that the encoded data follows a prior gaussian distribution, and thus combines the reconstruction loss with a distance function (between the gaussian prior and the actual learned distribution). For example, VAEs have been applied to scRNA-seq to predict cell response to biological perturbations [32]. Recently, [33], provided a supervised auto-encoder neural network that jointly predicts targets and inputs (reconstruction). However, neither VAEs [31] nor SAEs [33] provide a solution to the problem of relevant features and cells selections needed to increase the sensitivity of CRISPR-based perturbation associated with scRNAseq readout.

### SSAE criterion

In this section, we cope with these two issues by providing a sparse supervised autoencoder (SSAE) neural network method for selecting both relevant features (genes) and actual perturbed cells. **Figure 1** depicts the main constituent blocks of our proposed approach. Note that we added a “soft max” block to our SSAE to compute the classification score. Let *X* be the concatenated raw counts matrix (*n* × *d*) (n is the number of cells and d the number of genes) of control cells (targeted with a negative control gRNA) and gRNA-targeted cells for each target gene in a particular condition (Normoxia, Hypoxia 3h, 6h or 24h). Let *Y* be the vector of labels (*n* × 1) which component is 0 for control cells and 1 for the perturbed cell. Those labels, either “control” or “gRNA-targeted”, has been previously assigned for each cell according to the quantification of each gRNA of the CROP-seq library. Let Z be the encoded latent matrix (2 × 2). The matrix 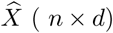 is the reconstructed data. *W* is the matrix of the weights of the the linear fully connected autoencoder neural network.

**Fig 1.**
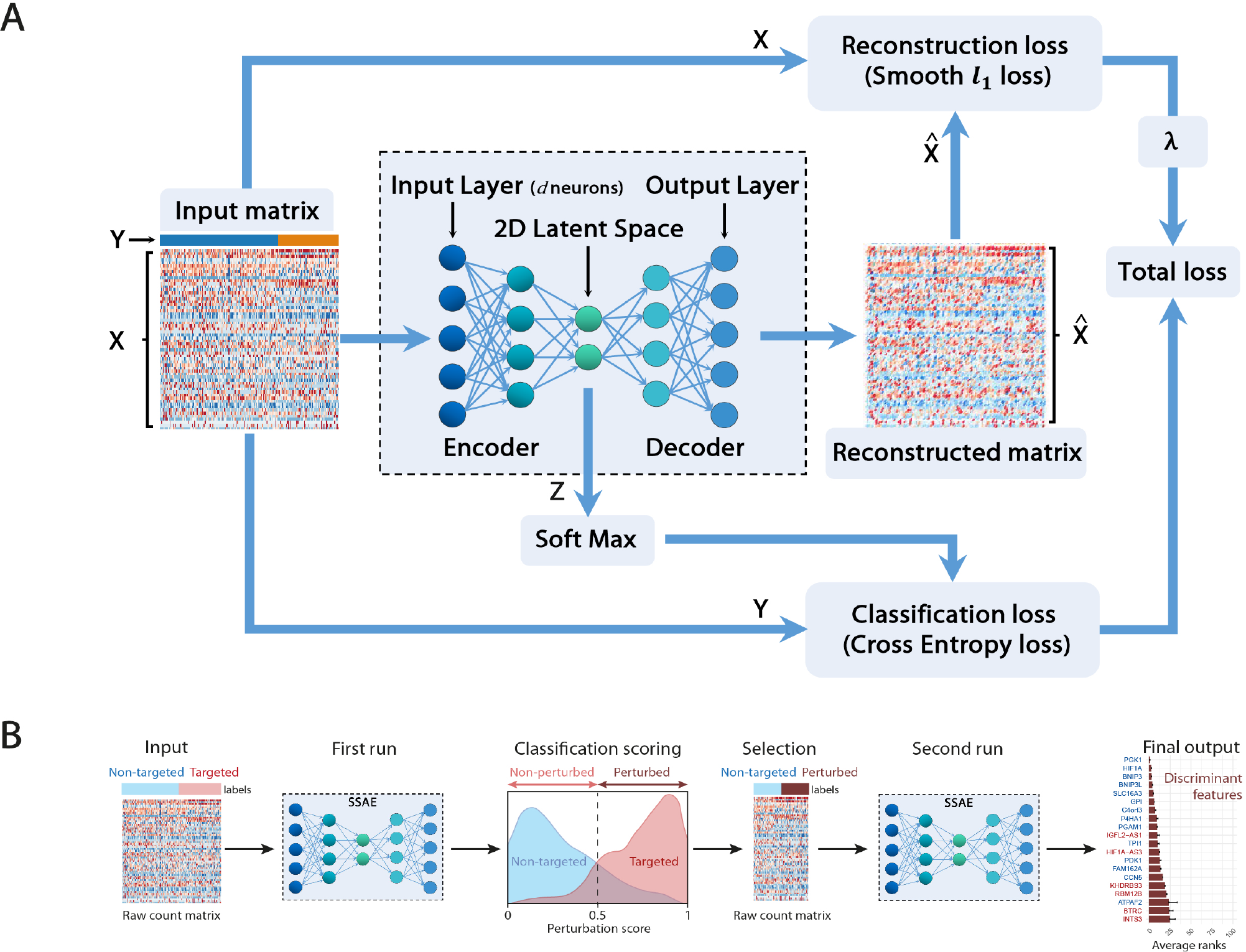
Sparse Supervised autoencoder (SSAE) framework: A: SSAE framework overview. B: Two-step SSAE classification of perturbed cells among gRNA-targeted cells.

The goal is to compute the network weights *W* minimizing the total loss which includes both the classification loss and the reconstruction loss. To perform feature selection, as large datasets often present a relatively small number of informative features, we also want to sparsify the network, following the work proposed in [34]. Thus, instead of the classical computationally expensive lagrangian regularization approach [35], we propose to minimize the following constrained approach [36]:

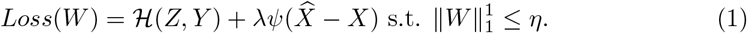

We use the Cross Entropy (CE) Loss for the classification loss ℋ. We use the robust Smooth *ℓ*_1_ (Huber) Loss [37] more robust than the mean square error (MSE) as the reconstruction loss *ψ*.

### Sparsity and gene selection using structured projections

A classical approach for structured sparsity is the Group LASSO method [38, 39] which consists of using the *ℓ*_2,1_ norm for the constraint on *W* . However, the *ℓ*_2,1_ norm does not induce an efficient sparse structured sparsity of the network [40], which leads to negative effects on performance.

In our method we achieve structured sparsity (feature selection) using the bilevel *ℓ*_1,1_ projection [34] of the weights W. We compute this bilevel *ℓ*_1,1_ projection using fast *ℓ*_1_ algorithms [41, 42]. We can also use the new *ℓ*_1,*∞*_ which provides similar sparsity performances [43]. Note that low values of *η* imply high sparsity of the network. We compute feature importance for the sparse supervised autoencoder using the SHAP method, implemented in the captum python package [44]. Those ranked weights give the top discriminating genes between the compared classes, which can be interpreted as the perturbation signature.

The main difference with the criterion proposed for VAEs in [31] and the criterion proposed for SAEs in [33] is the introduction of the constraint on the weights *W* to sparsify the neural network, in order to select relevant genes.

### Selecting actual perturbed cells using the softmax classifier

The goal of this section is to estimate the cells actually perturbed. We propose the following procedure thanks to the softmax formula. A first SSAE run gives a perturbation score thanks to the softmax layer [45] for both non-targeted control cells and for cells targeted for a particular gene.

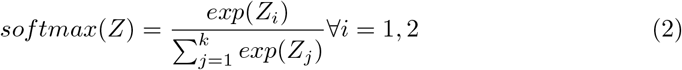

According to this specific score, called perturbation score, cells are separated into 2 subsets : targeted cells with a score *>* 0.5 are classified as “perturbed” cells, whereas targeted cells with a score *<* 0.5 are classified as “non-perturbed” cells. A new data matrix and a new label vector is generated, containing only the raw counts and labels of the selected perturbed cells and an equivalent number of randomly sampled non-targeted control cells in order to balance both classes. A second SSAE run provides a new list of the most discriminant features between both classes, ranked by their weight. This procedure is run multiple times with different initialization seeds in order to compute a mean and a standard deviation of the obtained ranks. The standard deviation ranks are used to evaluate the robustness of the perturbation signature. Again, neither VAEs [31] nor SAEs [33] provide a solution to the actual perturbed cell selection.

### Implementation of the SSAE framework

Following the work by Frankle and Carbin in [46], and further developed in [47], we follow a double descent algorithm, originally proposed as follows: after training a network, set all weights smaller than a given threshold to zero, rewind the rest of the weights to their initial configuration, and then retrain the network from this starting configuration while keeping the zero weights frozen. We replace the thresholding by our *ℓ*_1,1_ projection. We implemented our SSAE method using the PyTorch framework for the model, optimizer, schedulers and loss functions. We train the network using the classical Adam optimizer [48]. We used a symmetric linear fully connected network [22], with the encoder comprised of an input layer of *d* neurons, one hidden layer followed by a ReLU activation function and a latent layer of dimension *k* = 2 since we have two classes. The accuracy of the model, the mean and variance of the rank of selected genes was computed for each SSAE run using 4-fold cross-validation (which means that the train-validation split is random every time) and a mean over 3 seeds.

## 1 Results

### 1.1 Single-cell CRISPRi screening of hypoxia-regulated lncRNA

In order to gain new insights into the molecular functions of 6 hypoxia-regulated lncRNAs in LUAD cells we performed a single-cell CRISPRi transcriptome screening based on the CROP-Seq approach. We transduced A549 cells expressing double repressor Krab-MeCP2-dCas9 with a mini-library containing 12 validated gRNA targeting CYTOR (also known as LINC00152), LUCAT1, MALAT1, NEAT1, SNHG12 and SNHG21 as well as the two key regulators of the hypoxic response, HIF1A and HIF2/EPAS1. Each guide was individually validated by qPCR in A549 cells, showing a 75 to 95% inhibition of the target compared with control cells (**Table 1**). Two additional guides, with no effect on the genome, were used as negative controls. In order to mimic the hypoxic environment in which tumors develop in vivo, we equally divided the transduced dCas9-Krab-MeCP2 A549 cells in 4 samples that we then cultured in normoxia or in hypoxia during 3, 6 or 24 hours **Figure 2A**. Cells from each sample were labeled with a specific barcoded antibody (HTOs), pooled, and simultaneously sequenced using droplet based scRNA-seq (10X Genomics Chromium). The received gRNA and the culture condition were subsequently assigned for each cell by demultiplexing both gRNA and HTOs counts respectively.

**Fig 2.**
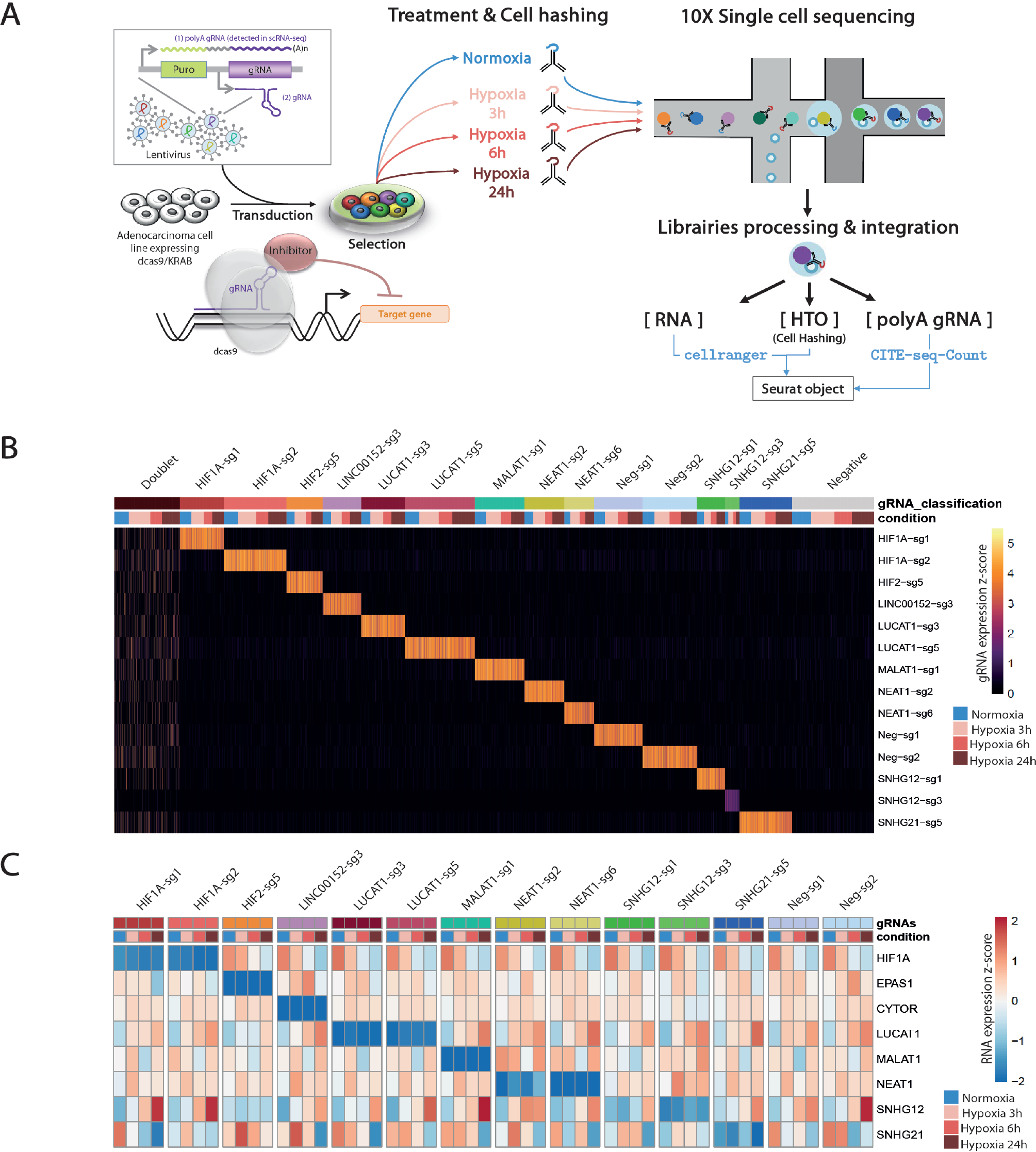
Single-cell CRISPRi screening: A: Design of CROP-seq experiment. B: Heatmaps of gRNA counts or target gene RNA in each cell, labelled according to assigned gRNA and condition after demultiplexing. C: Heatmap of target gene RNA in each cell, labelled according to assigned gRNA and condition after demultiplexing.

Overall, we found a balanced representation for each treatment and for each gRNA among the sequenced cells, except for the cells targeted by “SNHG12-sg3” which were depleted in all conditions (**Figure 2B, Table 2**). Moreover, the expression of this particular gRNA was lowly detected in those cells, confirming previous observations that this gRNA induced cell death and that only cells with low expression survive. Inhibition of target gene expression in the presence of their corresponding gRNA were validated in all 4 conditions, as well as their progressive increase (CYTOR, LUCAT1, NEAT1, SNHG12) or decrease (HIF1A and SNHG21) during hypoxia exposure (**Figure 2C**).

### 1.2 Mathematical and biological validation of the SSAE approach

In order to validate the SSAE approach, we first evaluated the effect of feature and cell selection on the accuracy of the model, and then the biological relevance of the detected transcriptomic perturbations induced by the knock-down of the two main regulators of the hypoxic response, HIF1A and HIF2.

#### 1.2.1 Feature and cell selection improve the accuracy of the model

We compared the performance of SSAE, for different criterion (MSE and Huber), with or without gene selection using *ℓ*_1,1_ constraint projection, and with or without cell selection on the dataset of HIF2-targeted cells and negative control cells cultured in hypoxic condition for 24h. Table 3 indicates that the SSAE with *ℓ*_1,1_ constraint projection and Huber criterion is able to discriminate both classes by selecting only a fraction of measured genes (19.46%), as it is shown in the matrix of connections between the first and second layer (**Figure 3A**). Moreover, the improvement of 5.39% of the model accuracy by using SSAE with *ℓ*_1,1_ constraint projection compared to SSAE without projection shows the efficiency of selecting only the most relevant features.

**Table 3.** Comparison of different criterion on HIF2 datase(CE is the cross entropy)

| Criterion | Gene selection (%). | F1 score (%). | accuracy (%). |
| --- | --- | --- | --- |
| CE + Huber and No-proj | 97 | 80.54 | 87.29 |
| CE + Huber + $\ell_{1,1}$ | 19.46 | 90.48 | 92.68 |
| CE + MSE + $\ell_{1,1}$ | 19.46 | 89.86 | 92.16 |

**Fig 3.**
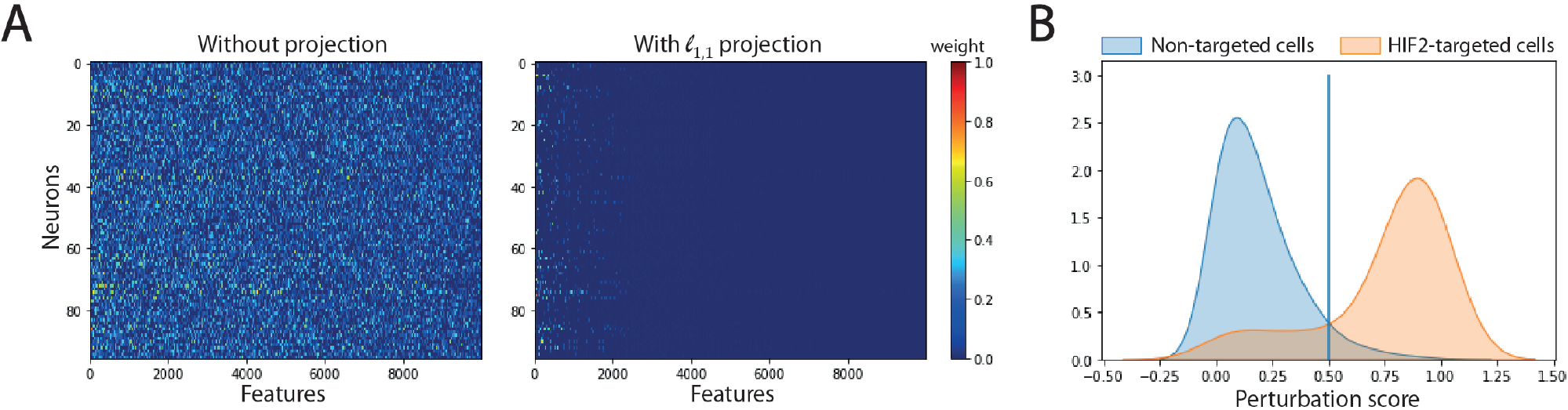
Impact of feature and cell selection on the accuracy of the model comparing HIF2-targeted cells and negative control cells after 24h of hypoxic exposure. A: Sparsity of the first layer: Left: using no projection, Right: using our *ℓ*_1,1_ projection. B: Distribution of the perturbation score for non-targeted and HIF2-targeted cells.

**Figure 3B** shows the distribution of perturbation scores computed with the softmax formula 2 for non-targeted and HIF2-targeted cells. Running the SSAE after selecting only perturbed HIF2-targeted cells, without feature selection, improves the accuracy by 4.67% (Table 4). Combining feature and cell selection improves the accuracy by 10.4%. Thus, we further analyzed the transcriptomic perturbations induced by the different gRNA using SSAE combining both gene and cell selection. The percentage of cells classified as perturbed among targeted cells for each gRNA in each condition are shown in **Figure 4**.

**Table 4.** SSAE Accuracy without or with feature and cell selection.

| SSAE (without feature and cell selection) | SSAE (only with feature selection) | SSAE (with feature and cell selection) |
| --- | --- | --- |
| 87.29% | 92.68% | 97.67% |

**Fig 4.**
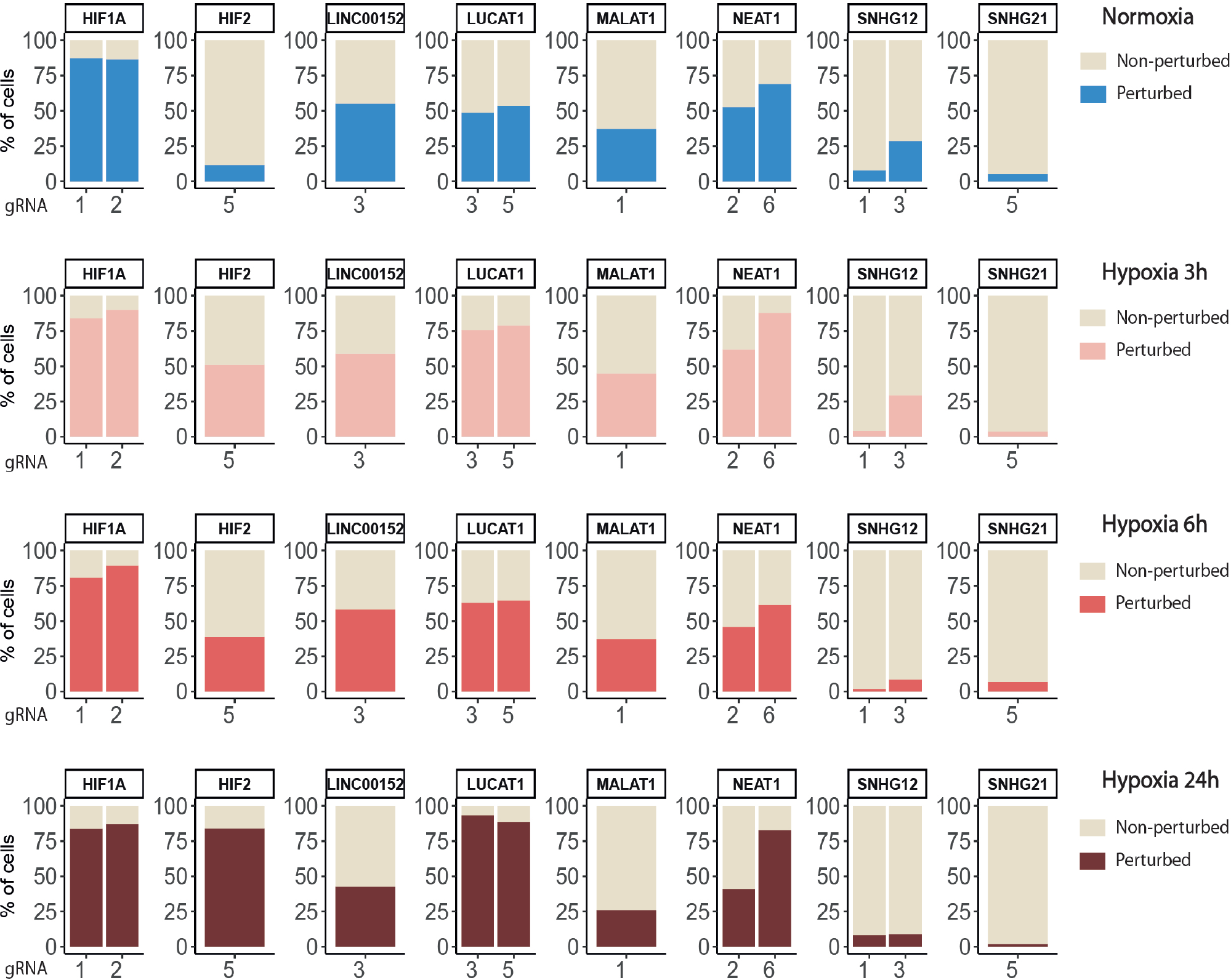
Percentages of targeted cells classified as perturbed or non-perturbed for each gRNA in each condition

#### 1.2.2 Knock-down of HIF1A and HIF2 differentially modulate the hypoxic response

Globally, the inhibition of HIF1A induced a strong transcriptomic perturbation which affected more than 85% of targeted cells in all conditions (**Table 5** and **Figure 4**).

**Table 5.** SSAE classification and accuracy for HIF1A and HIF2 gRNA targeted cells.

|  | Treatment | Targeted cells | Perturbed cells (%) | Accuracy (%) |
| --- | --- | --- | --- | --- |
| HIF1A | Normoxia | 475 | 86,7 | 95,33 |
| HIF1A | Hypoxia 3h | 554 | 87,5 | 94,00 |
| HIF1A | Hypoxia 6h | 372 | 85,8 | 93,67 |
| HIF1A | Hypoxia 24h | 554 | 85,7 | 94,67 |
| HIF2 | Normoxia | 147 | 11,6 | 66,67 |
| HIF2 | Hypoxia 3h | 202 | 51 | 86,33 |
| HIF2 | Hypoxia 6h | 104 | 38,5 | 82,33 |
| HIF2 | Hypoxia 24h | 213 | 84 | 97,67 |

Even in normoxic condition, the signature amplitude was sufficient to allow a classification accuracy above 93%. Among the genes modulated independently from the hypoxic status, we found SNAPC1, IGFL2-AS1, BNIP3L and LDHA, whereas PGK1, PDK1, or BNIP3 modulations were specific to hypoxic conditions (**Figure 5A**) . We also found gene modulations specific to early (KDM3A, HIPLDA, ZNF292, EGLN3) or late (SLC16A3, GPI, PGAM1, TPI1) hypoxic response, which correspond to the progressive establishment of the HIF1A-mediated metabolic switch [49]. In normoxia, the knock-down of HIF2 did not produce stable perturbations, except for its own target gene EPAS1 (**Figure 5B**). The 2 early time points of hypoxia exposure showed an improvement of the associated classification accuracy, which reflected a slight increase of the transcriptomic perturbation induced by HIF2 knock-down in these experimental settings. This early signature was mainly driven by genes involved in lipid metabolism ANGPTL4, IGFBP3 and HILPGA. Discrepancies between the results at 3h or 6h were mainly due to the lower number or targeted cells at 6h (104 instead of 202), which impacted the classification. At 24h of hypoxic exposure, the effect of HIF2 inhibition reached its maximum, with 84% perturbed cells and an accuracy of 97% (**Table 5** and **Figure 4**). However, this signature was quite different from that of HIF1A-targeted cells under the same condition. Indeed, some upregulated (ALDH3A1, CPLX2, FTL, PAPPA) or downregulated (ATP1B1, FXYD2, ANXA4, LOXL2) genes in HIF2-targeted cells were not modulated in HIF1A-targeted cells (**Figure 5C**). Moreover, several genes showed an opposite perturbation between the two groups of cells. This was the case for BNIP3, PGK1, GPI, FAM162A, SLC16A3, TPI1, or PGAM1 which were downregulated upon HIF1A inhibition but were found upregulated upon HIF2 inhibition after 24h of culture in hypoxia. These results were consistent with the known role of HIF2, which is activated upon prolonged exposure to hypoxia and is involved in the regulation of the chronic hypoxic response [5]. They also confirm that in LUAD cells, HIF1A and HIF2-regulated functions are specific, or even antagonistic for certain genes, which has been previously demonstrated in other cancers [50].

**Fig 5.**
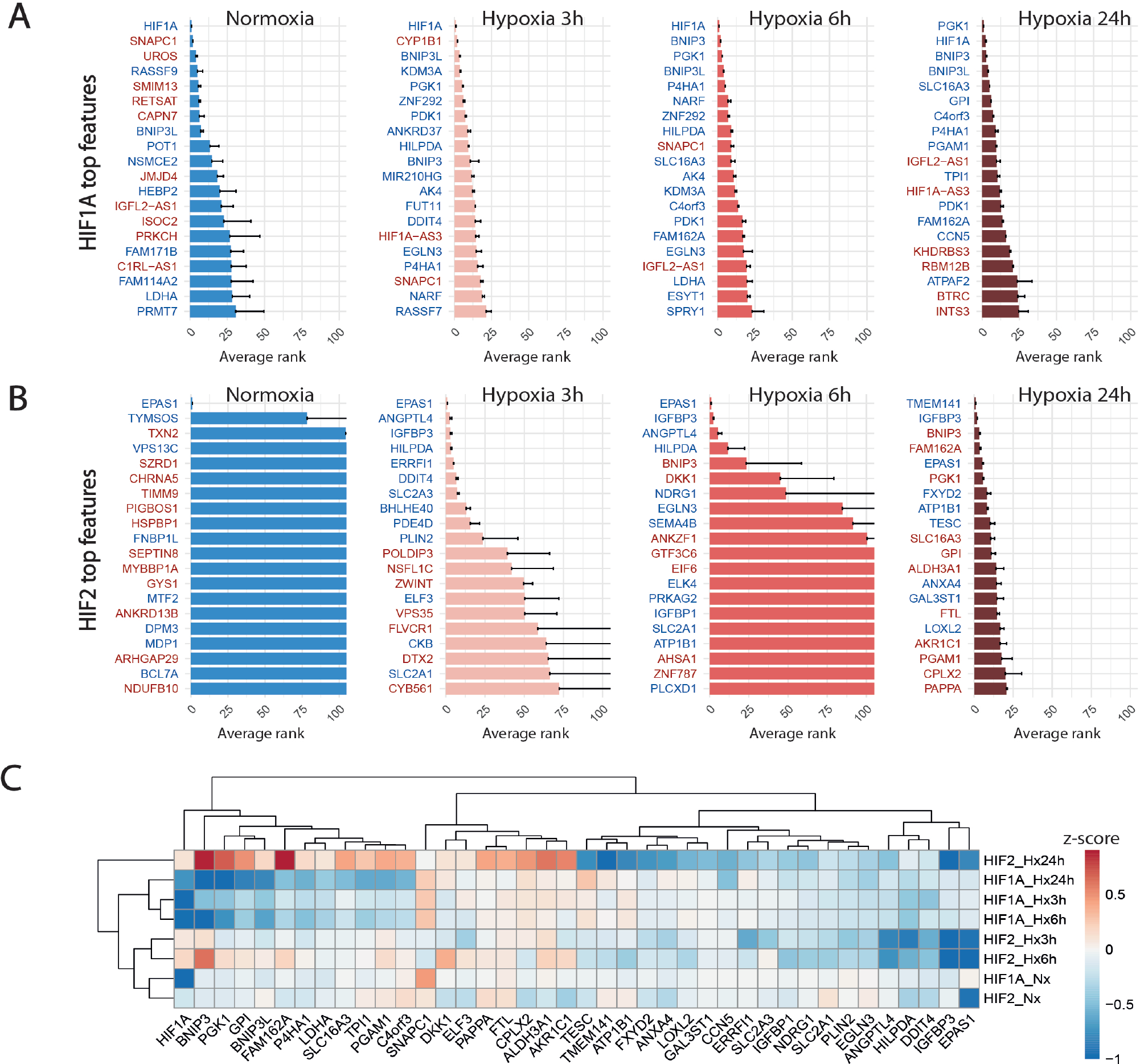
Knock-down of HIF1A and HIF2 differentially modulate the hypoxic response. A: Top 20 discriminant features between perturbed and control cells for HIF1A for each treatment. Upregulated or downregulated genes are written in red or blue respectively. B: Top 20 discriminant features between perturbed and control cells for HIF2/EPAS1 for each treatment. C: Differentially expressed genes between perturbed and control cells for HIF1A and HIF2 for each treatment.

### 1.3 Knock-down of hypoxia-regulated lncRNA LUCAT1 leads to hypoxic condition-dependent transcriptomic modulations

We then applied the SSAE method to classify cells treated with the 6 gRNA targeting hypoxia-regulated lncRNAs and cultured in the 4 conditions. Globally, the SSAE was able to classify perturbed and control cells with a good overall accuracy around 80%, except for SNHG12 and SNHG21 (**Table 6** and **Figure 4**). However, despite their promising accuracies, we did not detect any other stable perturbations than the target gene for both MALAT1 and NEAT1 targeted cells, as indicated by the obtained high means and standard deviations of the computed ranks, while those two lncRNAs were previously associated with various gene regulation functions [51, 52] (**Figure 6A-B**). The SSAE outcomes were different for the classification of LUCAT1-targeted cells. Indeed, the transcriptomic inhibition of LUCAT1 resulted in a stable upregulation for PCOLCE2 and ISCA1 in normoxia, HDHD2 after 3h, and 6h of hypoxia (**Figure 6C**). ISCA1 and HDHD2 encode for metal ion binding proteins, whereas TFCP2 is a known oncogene. After 24h of hypoxia, a completely different perturbation signature was found, with at least 6 stably modulated genes, including the upregulation of KDM5C, TMEM175 and NIT1, as well as the downregulation of ATP6AP1, PEX1 and PHF20. ATP6AP1 and PEX1 are respectively components of the V-ATPase and the peroxisomal ATPase complexes, while TMEM175 is a proton channel also involved in pH regulation. KDMC5 and PHF20 are both involved in chromatin remodeling and transcriptomic regulation, while NIT1 is associated to tumor suppressor functions. This particular signature allowed the classification of 90,4% of targeted cells with an accuracy of 85,67% (**Table 6** and **Figure 4**). These results indicate that LUCAT1 inhibition may induce hypoxic condition-dependent transcriptomic modulations that potentially impact tumor survival and gene regulatory processes during prolonged exposure to hypoxic conditions, completing our previous observations [12].

**Table 6.** Supervised autoencoder classification and accuracy for hypoxia regulated lncRNAs gRNA targeted cells (*obtained without cell selection)

|  | Treatment | Targeted cells | Perturbed cells (%) | Accuracy (%) |
| --- | --- | --- | --- | --- |
| LUCAT1 | Normoxia | 438 | 51,8 | 73,33 |
| LUCAT1 | Hypoxia 3h | 583 | 77,7 | 82,33 |
| LUCAT1 | Hypoxia 6h | 391 | 63,9 | 79,00 |
| LUCAT1 | Hypoxia 24h | 666 | 90,4 | 85,67 |
| MALAT1 | Normoxia | 205 | 37,1 | 82,67 |
| MALAT1 | Hypoxia 3h | 269 | 45 | 82,33 |
| MALAT1 | Hypoxia 6h | 194 | 37,1 | 78,00 |
| MALAT1 | Hypoxia 24h | 241 | 26,1 | 78,67 |
| NEAT1 | Normoxia | 274 | 59,1 | 86,33 |
| NEAT1 | Hypoxia 3h | 390 | 73,3 | 89,00 |
| NEAT1 | Hypoxia 6h | 237 | 52,7 | 85,33 |
| NEAT1 | Hypoxia 24h | 566 | 58,7 | 87,33 |
| LINC00152 | Normoxia | 147 | 55,1 | 87,67 |
| LINC00152 | Hypoxia 3h | 200 | 59 | 91,67 |
| LINC00152 | Hypoxia 6h | 150 | 58 | 86,33 |
| LINC00152 | Hypoxia 24h | 209 | 42,6 | 85,00 |
| SNHG12 | Normoxia | 196 | 14,8 | 65* |
| SNHG12 | Hypoxia 3h | 217 | 14,7 | 69* |
| SNHG12 | Hypoxia 6h | 154 | 3,9 | 67,4* |
| SNHG12 | Hypoxia 24h | 213 | 8,5 | 71* |
| SNHG21 | Normoxia | 211 | 5,2 | 60.7* |
| SNHG21 | Hypoxia 3h | 256 | 3,5 | 61.1* |
| SNHG21 | Hypoxia 6h | 192 | 6,8 | 59,1* |
| SNHG21 | Hypoxia 24h | 299 | 2 | 63,5* |

**Fig 6.**
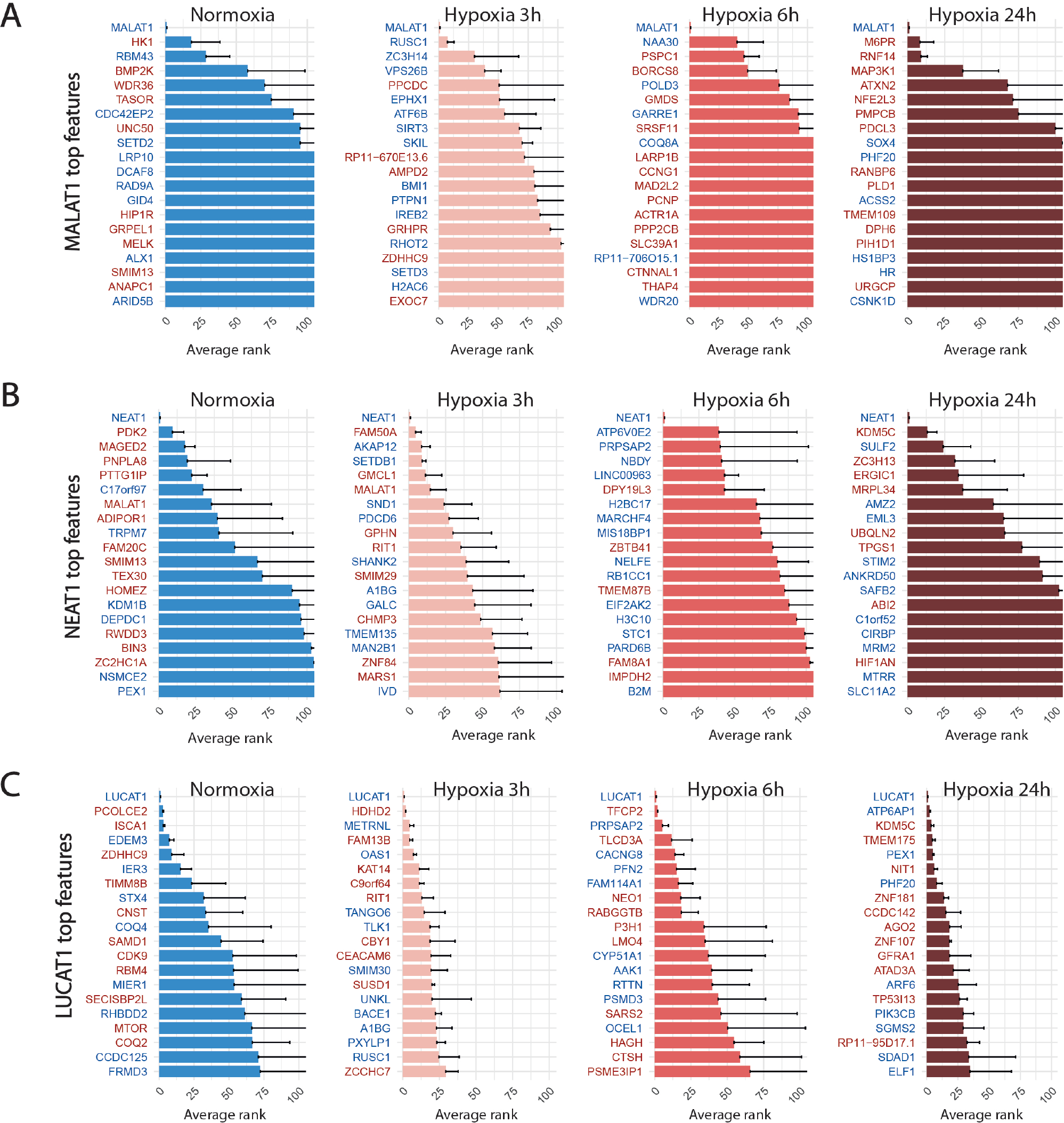
Top 20 discriminant features between perturbed and control cells for LUCAT1 (A), MALAT1 (B) and NEAT1 (C) for each treatment. Upregulated or downregulated genes are written in red or blue respectively.

For LINC00152, the combined inhibition of CYTOR/LINC00152 with MIR4435-2HG (**Figure 7A**), whose sequences are highly homologous (99% in the 220 bp region including the most efficient gRNA), was sufficient to select half of the targeted cells with an accuracy above 85% regardless of the condition (**Table 5** and **Figure 4**).

**Fig 7.**
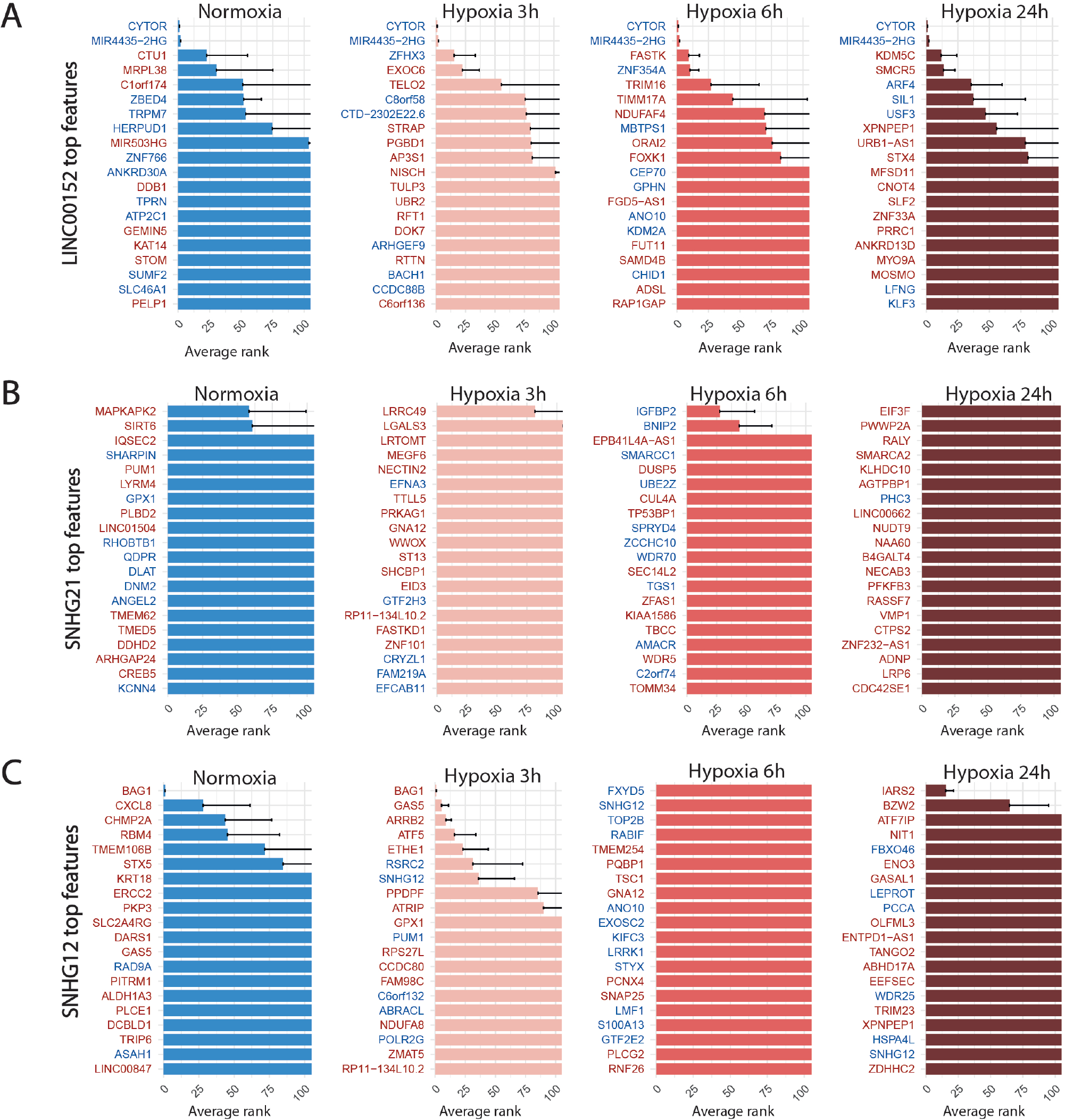
Top 20 discriminant features between perturbed and control cells for LINC00152 (A), SNHG21 (B) and SNHG12 (C) for each treatment. Upregulated or downregulated genes are written in red or blue respectively.

For the SNHG12 and SNHG21 datasets, the first round of SAE selected only around 10% of perturbed cells (**Table 6** and **Figure 4**). Thus we could not run the SAE for the second round because of a too low number of cells for the 4 fold cross validation. For those 2 genes, we just reported the average accuracies obtained after the first round of the SSAE (**Table 6**). As SNHG21 expression is relatively low in LUAD cells and is decreased by hypoxic stress, the extent of its inhibition was therefore weaker and not sufficient to distinguish targeted from control cells. Combined with the lack of transcriptomic effect induced by its knock-down, it explains the poor classification results and the randomness of features selected for cells targeted by this particular gene under all conditions (**Figure 7B**).

### 1.4 SSAE classification revealed an anti-apoptotic signature expressed by a subset of SNHG12-targeted cells in response to the cytotoxic effect of one of its gRNA

Looking at the SSAE classification outcomes for SNHG12-targeted cells, only about 15% of them were classified as perturbed in normoxia and after 3h of hypoxia, with a poor accuracy (**Table 6** and **Figure 4**). The number of selected cells was even worse for a longer exposure to hypoxia.

Nevertheless, the ranked list of top discriminant features between the few perturbed cells and control cells obtained for the first two time points showed a notable perturbation signature. In normoxia, it was only composed of BAG1 upregulation, whereas after 3h of hypoxia exposure, this signature was completed by GAS5 (snoRNAs-containing lncRNA gene), ARRB2, ATF5, and ETHE1 upregulations (**Figure 7C**). These 5 genes are all known anti-apoptotic factors. We hypothesized that this anti-apoptotic signature was expressed by a subset of LUAD cells that were actively escaping the cytotoxic effect we systematically observed for the most efficient of the two gRNA selected for targeting SNHG12, SNHG12-sg3 (**Table 1**). Indeed, most of the cells classified as perturbed were specific to this particular gRNA (**Figure 4**). As this signature progressively attenuated over time under hypoxic conditions, we speculate that the activation of this anti-apoptotic response may be inhibited by hypoxic stress, or that hypoxia may protect against the cytotoxic effect of this guide. These results demonstrate the precision of the SAE-based approach to detect a short signature, even restricted to a small subset of cells.

### 1.5 Comparison of SSAE with others machine learning methods

We compared the classification performance and the biological relevance of extracted features between the SSAE, with and without cell selection of the most responsive cells, SAE [33] and Random Forests using 400 estimators and the Gini importance (GI) for feature ranking. We performed this comparison for 2 representative datasets, namely HIF2-targeted cells versus control cells and LUCAT1-targeted cells versus control cells following 24h of hypoxia exposure.

For the first dataset, as HIF2 inhibition induced a strong perturbation signature, the 10 first selected features between all methods were highly similar, even between SSAE/SAE and Random Forest (**Figure 8**A). However, using the SSAE with cell selection outperforms Random Forest with an increase of 25.25% of accuracy.

**Fig 8.**
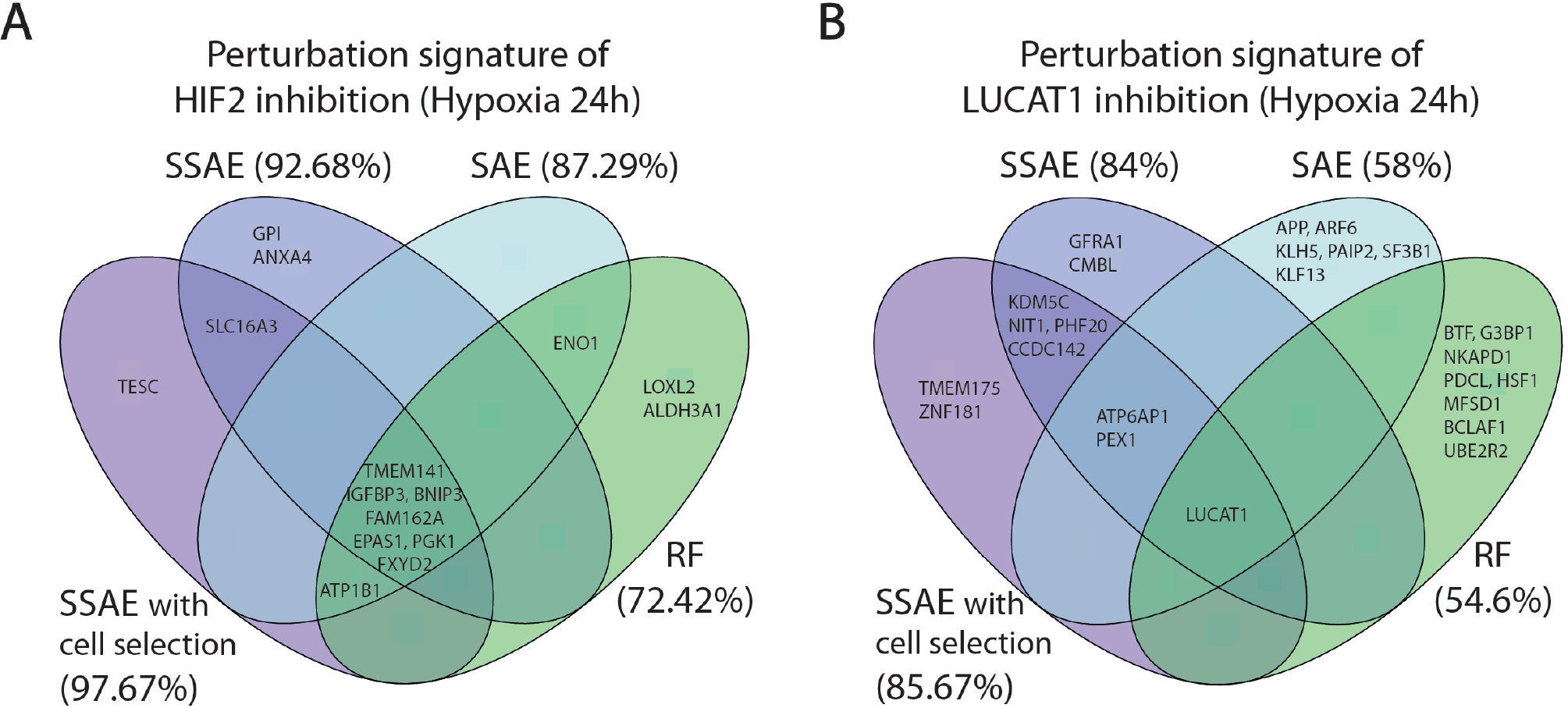
Comparison of the 10 first selected features between SSAE (with or without cell selection), SAE and Random Forests, for HIF2/EPAS1 (A) or LUCAT1 (B) targeted cells in hypoxia 24h. Associated accuracies are indicated in brackets.

For LUCAT1 dataset, few overlaps were found between the first 10 selected features with each method. The inhibition of the target gene LUCAT1 was the only feature commonly detected (**Figure 8B**). ATP6AP1 and PEX1 were the only two overlapping genes between SSAE and SAE obtained signatures, while KDMC5, NIT1, PHF20 and CCDC142 were specific to SSAE regardless of the cell selection step.

While the signature of Random Forests was very specific, note that the author of RF proposes two measures for feature ranking, the variable importance (VI) and Gini importance (GI): [53] showed that if predictors are real with multimodal Gaussian distributions, both measures are biased. Moreover, since using the SSAE with cell selection outperforms RF by 31.07 %, SAE by 27% and SSAE without cell selection by 1,67%, it is reasonable to claim that the SSAE perturbation signature is the most relevant.

## Discussion

Single-cell CRISPR(i)-based transcriptome screenings are powerful tools for simultaneously accessing the expression profiles of cells targeted by different gRNA, in order to infer target genes functions from the observed perturbations. However, these approaches are limited by the low molecule capture rate and sequencing depth provided by droplet-based scRNA-seq, which produce sparse and noisy data. Furthermore, the outcome of CRISPR-induced modification in each cell is a stochastic event, depending among other things, on the expression levels of the transcribed gRNA and dCas9, as well as the accessibility of the target gene locus, that may be heterogeneously regulated at the epigenomic levels in the different cells. For these reasons, the induced perturbation signature and its detection are likely heterogeneous between cells, even when dCas9-expressing cells receiving the same gRNA have been cloned. Deciphering this heterogeneity in sparse data is even more complex when the targeted genes are not master genes involved in signaling or regulatory pathways, such as transcription factors and receptors. In this respect, a previous study [20] has shown that this particular challenge cannot be met using conventional scRNA-seq analysis tools such as differential expression, which is clearly limited to the detection of weak and heterogeneous perturbation signals. This challenge seems even more complex for the study of perturbations mediated by knockdown of non-coding RNAs, which have been largely involved in the fine-tuning of gene expression regulation. To increase the sensitivity of single-cell CRISPR(i)-based transcriptome screenings, we propose here a powerful feature selection and classification approach based on a sparse supervised autoencoder (SSAE). It leverages in particular on the known cell labels initially given by gRNA counts demultiplexing to constrain the latent space to fit the original data distribution. Beyond high statistical accuracy, our SSAE offered relevant properties that distinguishes it from classical classification methods : i) a stringent feature selection producing an interpretable readout of ranked top discriminant genes associated to their weights; ii) a classification score which allow the selection of the most perturbed cells and the eventual signal to obtain a more robust perturbation signature. We first validated this approach by analyzing the perturbations associated with the knock-down of the two master regulators of the hypoxic response, HIF1A and HIF2. We showed that the SSAE was able to learn a latent space and a perturbation signature which can for exemple almost perfectly discriminate HIF2-targeted cells from their control in condition of prolonged hypoxia. The SSAE classification accuracy provided a global perturbation score associated with HIF1A and HIF2 at each time point, reflecting the biological activity of each factor during the hypoxic response. We were able to recapitulate the known distinct influence and target specificity of HIF1 and HIF2 during the hypoxia time course [5], with notably i) a strong perturbation driven by HIF1 at early time points; ii) a progressive influence of HIF2 with a maximum effect observed at 24h of hypoxia; iii) a specificity regarding their targets, with sometimes an opposite regulation for some genes. Finally, this unique dataset provides a global and dynamic description of the transcriptomic modulations mediated by the two main regulators of the hypoxic response in LUAD A549 cells. Surprisingly, we did not detect any relevant and stable perturbation in cells targeted for LINC00152, MALAT1, NEAT1 and SNHG21, in the four culture conditions. This result appears quite unexpected for MALAT1 and NEAT1, two of the most studied lncRNAs that are associated with various functions in cancer, including proliferation, migration, and invasion [51, 54]. In particular, it has been shown that MALAT1 knockout in the same cellular model (A549) modulated a set of metastasis-associated genes [55]. Although CRISPRi-mediated knock-down achieved an efficient knock-down (*>* 95%), it is however possible that based on the very high level of MALAT1, the remaining transcripts are sufficient to mediate the cellular function. Another possibility could be due to differences in methodology, notably the need to isolate single clones for the knockout protocol, a long procedure that can profoundly affect the transcriptome, compared with the CROP-seq approach performed on a bulk population prior to immediate single-cell isolation. A similar situation may occur for NEAT1, a highly abundant lncRNA acting as a structural scaffold of membraneless paraspeckle nuclear bodies. Moreover, NEAT1 can produces two isoforms, with a differential regulation upon stress and distinct functions [56].Additional work will be thus necessary to further analyze the relative proportion of the two isoforms in A549 cells and their potential function during hypoxia.

However, for LUCAT1-targeted cells after 24h of hypoxia exposure, we found a stable signature of 6 modulated genes, which are associated with pH or gene regulation. It suggested a potential capacity of LUCAT1 to promote tumor cell survival during prolonged hypoxia and to contribute to an aggressive phenotype in LUAD cells, as we previously demonstrated [12]. Finally, we also found a relevant signature in SNHG12-targeted cells, characterized by the upregulation of anti-apoptotic genes. As this signature is almost exclusive to cells targeted by the most effective gRNA against SNHG-12, which appeared to systematically induce cell death, we hypothesized that it is expressed by surviving cells. The potential pro-oncogenic role of the complex SNHG-12 locus, producing a lncRNA and 3 snoRNAs, should be pursued to decipher the molecular components associated with this phenotype, as also suggested by previous studies [57].

In this paper, we demonstrate that the SSAE is highly relevant in situations in which low signals in a restricted number of cells need to be detected. However, performances (accuracy, F1 score, AUC …) of the SSAE (similarly to all statistical method) are highly dependent on the number of samples/cells compared. Low cell number impact classification performance and can produce inconsistent results, such as better accuracy and robustness of selected features for HIF2-targeted cells after 3h of hypoxia compared with 6h exposure. In this context, the relevance of the top selected genes list and their superiority over other compared methods can be asserted by evaluating the robustness of the ranks and the classification accuracies. The size of the perturbation signatures obtained for LUCAT1 and SNHG12 datasets prevented the utilization of functional enrichment analysis to characterize their modulated functions. Moreover, as these small signatures were found in specific subsets of targeted cells and dynamically during the hypoxic response, it appears very difficult to validate them using a global experimental approach that will average the signal across all cells. Despite these limitations, we believe that our approach is well suited to the particular deciphering of single cell CRISPR-based screen with omics readout, or for other similar assays to assess the effect of perturbation at the single cell level.

## Availability of data and materials

We implemented the SSAE code with python. Associated functions and scripts, as well as all input matrices used in the study are available at https://github.com/MichelBarlaud/SAE-Supervised-Autoencoder-Omics. Raw sequencing files and counts matrices of total UMI, HTOs, and gRNA will be deposited in the Gene Expression Omnibus. The scripts used for data processing and analysis will be available on github (https://github.com/marintruchi) by the time of publication.

## Competing interests

The authors declare that they have no competing interests.

## Fundings

We acknowledge the support from the Centre National de la Recherche Scientifique (CNRS), Université Côte d’Azur, Canceropôle PACA (Action Structurante CRISPR SCREEN), the French Government (National Research Agency, ANR) program “Investissements d’Avenir” UCAJEDI n° ANR-15-IDEX-01 (PERTURB-ENCODER) and ANR-22-CE17-0046-01 MIR-ASO, Plan Cancer 2018 “ARN non-codants en cancérologie : du fondamental au translationnel” (number 18CN045) and Fondation ARC.

## Acknowledgments

The authors thank the technical support of the UCA GenomiX of the University Côte d’Azur. We also thank A. Monteil and C. Lemmers from the Vectorology facility, PVM, Biocampus Montpellier, CNRS UMS3426

## Notes

https://github.com/MichelBarlaud/SAE-Supervised-Autoencoder-Omics

### Competing Interest Statement

The authors have declared no competing interest.

### Summary of Updates

- Providing more details about the SSAE approach (see materials and Methods, new section "2.7 Method: a new sparse supervised autoencoder neural network (SSAE)" - Evaluation of the SSAE method with or without selection, see new tables 3-4 and new Figure 3. - Benchmarking of the SSAE method against a Supervised autoencoders (SAE), see new Fig.8 - new presentation of the data presented in new Figures 5, 6 and 7 - Major modifications of the material and method as well as the results sections - Minor modifications of the abstract, introduction and discussion

